# Monocarboxylate Transporters are Involved in Extracellular Matrix Remodelling in Pancreatic Ductal Adenocarcinoma

**DOI:** 10.1101/2022.02.01.478448

**Authors:** Ayşe Ufuk, Terence Garner, Adam Stevens, Ayşe Latif

## Abstract

Pancreatic ductal adenocarcinoma (PDAC) is an aggressive malignancy with a five-year survival rate of <8%. PDAC is characterised by desmoplasia with abundant extracellular matrix (ECM) rendering current therapies ineffective. Monocarboxylate transporters (MCTs) are key regulators of cellular metabolism and are upregulated in different cancers, however their role in PDAC desmoplasia is little understood. Here, we investigated MCT and ECM gene expression in primary PDAC patient biopsies using RNA-sequencing data obtained from Gene Expression Omnibus. We generated a hypernetwork model from these data to investigate whether a causal relationship exists between MCTs and ECMs. Our analysis of stromal and epithelial tissues (n=189) revealed 9 differentially expressed MCTs, including upregulation of SLC16A2/6/10 and the non-coding SLC16A1-AS1, and 502 ECMs including collagens, laminins, and ECM remodelling enzymes (false discovery rate<0.05). Causal hypernetwork analysis demonstrated a bidirectional relationship between MCTs and ECMs; 4 MCT and 255 ECM-related transcripts correlated with 90% of differentially expressed ECMs (n=376) and MCTs (n=7), respectively. The hypernetwork model was robust, established by two independent approaches involving iterated sampling and silencing of indirect interactions in the network. This transcriptomic analysis highlights the role of MCTs in PDAC desmoplasia via associations with ECMs, opening novel treatment pathways to improve patient survival.

**Simple Summary:** Monocarboxylate transporters (MCTs) carry a variety of substrates with MCT1-4 being well characterised and involved in proton-coupled transport of monocarboxylates (such as lactate) which can be used as metabolic fuel for cancer cells. Increased acidity of tumour microenvironment via MCTs favours remodelling of extracellular matrix (ECM) leading to desmoplasia associated with tumour metastasis and poor patient outcomes. Although MCT1-2/4 are upregulated in several cancers, their expression and role in pancreatic ductal adenocarcinoma desmoplasia is little understood. Here, we aimed to understand the role of MCTs in desmoplasia through their associations with ECM components. Our analysis using hypernetworks showed the presence of bidirectional associations of MCTs and ECMs, suggesting the presence of a causal relationship and the need to further investigate their functional associations. It confirms the role of MCTs in desmoplasia highlighting their importance as therapeutic targets alone or in combination with key ECM components to potentially improve patient outcomes.

## 1. Introduction

Pancreatic cancer is the 12^th^ most common cancer and the 7^th^ highest cause of cancer mortality worldwide [1]. In the UK, it is the 10^th^ most common malignancy with its incidence rates predicted to increase by 6% by 2035 [2]. PDAC is the most common malignancy of the pancreas accounting for 95% of all pancreatic cancer cases with five-year survival rate of less than 8% [3, 4]. Poor clinical prognosis of PDAC stems from the difficulty of its diagnosis, ineffective treatment options at advanced stages, and surgery being the only existing curative option that can be offered to only 10-20% of patients with resectable PDAC [4-6].

A key feature of PDAC is the existence of extensive fibrotic stroma, also known as desmoplasia, which has an abundance of dense extracellular matrix (ECM) surrounding fibroblasts, immune cells, endothelial cells, and neuronal cells [7, 8]. The components of ECM include collagens, glycoproteins (e.g., laminins, elastin, and fibronectin), proteoglycans (e.g., hyaluronic acid) as well as ECM remodelling enzymes involved in degradation or cross-linking of the ECM components (e.g., matrix metalloproteinases (MMPs) and their tissue inhibitors (TIMPs) and lysl oxidases (LOXs)) [9-11]. The excessive accumulation of ECM components has been highlighted as a major contributor to PDAC progression and the resistance to therapeutic efficacy [5, 9]. ECM remodelling occurs as a result of the crosstalk between tumour cells and the microenvironment including pancreatic satellite cells in stroma which cause aberrant secretion of ECM components by the secretion of pro-fibrotic and inflammatory growth factors.

Monocarboxylate transporters (MCTs) are key players in cellular metabolism and have an important role in regulating intracellular pH [12]. They belong to the solute carrier 16 (SLC16) gene family with MCT1/*SLC16A1*, MCT2/*SLC16A7*, MCT3/*SLC16A8*, and MCT4/*SLC16A3* being well characterised in terms of their role in proton coupled transport of monocarboxylates such as lactate, pyruvate, and ketone bodies. From metabolic standpoint, most solid tumours rely on glycolysis to produce energy leading to the production of large amounts of lactate that is exported by the MCTs [13, 14]. Therefore, MCTs help cancer cells maintain their high glycolytic activities and contribute to tumour acidosis and progression [15, 16]. MCT1, MCT2, and MCT4 are known to be upregulated in different cancers and high expression of MCT1 and MCT4 is often associated with poor prognosis [13, 15]. MCTs contribute to the acidification of tumour microenvironment which is known to activate and increase the expression of ECM remodelling enzymes involved in ECM degradation, contributing to desmoplastic reaction [17, 18]. However, their expression and association with desmoplasia is still largely unknown in PDAC.

So far, the association between MCTs and ECM components have been shown in a limited number of cancer types *in vitro*. For example, upregulation of MCT1 was associated with increased invasiveness and migration of nasopharyngeal carcinoma cell lines accompanied with increased expression of MMP2 and MMP9 and downregulation of TIMP1 and TIMP2 [19]. Secondly, overexpression of lysyl oxidase like 1 (LOXL-1) in a non-small cell lung cancer metastasis model was associated with upregulation of MCT1/2, increased expression and activity of MMP2/9, increased/enhanced cell migration and invasion along with increased extracellular lactate at a low pH [20]. The latter findings also support correlation of acidic pH via lactate transport by the MCTs with increased activity of the MMPs [18, 21, 22]. However, in PDAC, the link between MCT and ECM components have not been elucidated. Therefore, it is vital to investigate the correlation of MCTs with ECMs and consolidate the role of MCTs in PDAC desmoplasia as well as bringing forward novel treatment strategies co-targeting MCTs and key ECM players to reduce tumour progression and improve patient survival.

‘Omics approaches are now widely used in the field of cancer to identify dysregulated molecular mechanisms, provide insights into biological pathways, and propose potential biomarkers for different cancer types. Incorporation of higher-order interactions driven from ‘omic data is now seen as an essential part of modelling biological systems [23]. Hypernetworks model higher-order interactions or relationships between ‘omic elements (represented by nodes) based on large numbers of shared correlations (represented by edges) [24]. Such interactions which are normally not captured by traditional pairwise transcriptomic approaches provide a model of functional relationships between these ‘omic elements [25, 26]. In this study, we set out to evaluate MCT-ECM interaction using *in silico* approaches utilizing data available in public repository, the National for Biotechnology Information (NCBI) Gene Expression Omnibus (GEO). To achieve this, we investigated the causal link between MCT and ECM interaction using a hypernetwork model of PDAC transcriptome based on literature collated RNA-sequencing data. In order to refine the hypernetwork model, differential expression of MCT and ECM genes were first determined. In addition, we sought to understand the correlation between MCT and ECM expression with age at PDAC diagnosis. Finally, we assessed MCT and ECM expression in shortand long-term survivors of patients with PDAC.

## 2. Materials and Methods

### Study selection and data processing

The NCBI GEO repository browser (https://www.ncbi.nlm.nih.gov/geo/browse/) was used to search for RNA-sequencing (RNA-seq) studies of pancreatic cancer involving “pancreatic” as a search term, “homo sapiens” as an organism, and “expression profiling by high throughput sequencing” as a series type. Studies with RNA-seq data from primary PDAC tissue biopsies with their matching controls where available were included in the analysis.

The quality control for the raw sequencing data was performed with FastQC v0.11.9 (https://www.bioin-formatics.babraham.ac.uk/projects/fastqc/). The raw sequencing reads were filtered using BBDuk from the BBMap toolkit v38.87 (ktrim=r, k=21, mink=9, hdist=1; https://sourceforge.net/projects/bbmap/) to remove adapters and quality-trim both ends to Q15 (qtrim=rl, trimq=15, minlength=36). Read mapping to human reference genome hg38 with Gencode v35 annotations and gene quantification using “--quantMode GeneCounts” were performed with STAR v2.7.6a [27]. Post-mapping quality control was performed with RseQC v4.0.0 (http://rseqc.sourceforge.net/).

### Clustering analysis

To evaluate the quality of the collated transcriptomic data and assess the similarities of different datasets, unsupervised clustering analysis was performed using Principal Component Analysis (PCA). For the analysis of all collated studies (n=4), each study was initially processed independently to remove genes with low expression (genes with fewer than 10 reads) using the R package edgeR (version 3.32.1; [28]). Once genes with low expression were filtered out, the raw count table including all datasets was trimmed down to these common genes. Trimmed Mean of M-values (TMM) normalisation was used to facilitate comparison of expression between samples. These normalised data were then used to conduct PCA using the R package mixOmics (version 6.14.1, [29]). The analysis was run with the pca function based on the calculation of the first two principal components while both centring and scaling of the data were applied.

Clustering of the stromal and epithelial dataset alone (GSE93326) based on tumour overall stage, grade, and N-score was also performed by applying each of these features as a factor in the analysis.

### Differential gene expression analysis

Differentially expressed MCT, ECM and ECM-related genes were identified using the R package edgeR. When analysing differentially expressed genes (DEGs) in the stromal and epithelial dataset alone (GSE93326), the comparison was performed between stroma and epithelium, which included 66 matched tissues and additional 57 stromal data in comparison to the analysis conducted by Maurer *et al*. (2019) (analysed 60 matched tissues) [30]. Analysis of DEGs in short-(STS) and long-term survivors (LTS) from the GSE79668 dataset was performed using tumour tissues from the mentioned survival groups (14 and 13 samples from STS and LTS, respectively). When analysing DEGs between tumour and non-tumour tissues from all datasets combined (n=4), the pre-processing of the studies in terms of removal of low expression genes and data normalisation was handled as described above. Gene expression levels were calculated as log2 count per million (CPM). The difference in gene expression levels was calculated as the log2-fold changes (logFC) of genes between the comparison groups. Genes were filtered using a false discovery rate (FDR)-adjusted p-value <0.05 to determine the significance. Hierarchical clustering analysis of the DEGs was done by complete method with Euclidean distance. The gene expression profiles were visualised with heatmaps using a modified version of the heatmap.2 function of the R package gplots (version 3.1.1) [31] to allow simultaneous visualization of multiple annotations. Log-normalised expression levels of the MCT genes were visualised with violin plots using the R package ggplot2 (version 3.3.5) [32].

We identified differentially expressed ECM and ECM-related genes based on the NABA_MATRISOME gene set (version 5.0) from Molecular Signature Database (MSigDB) as a reference [10]. This gene set is an ensemble of 1026 genes which encode ECM and ECM-associated proteins. Due to their roles in cell adhesion and migration, and potential contributions to ECM remodelling and stiffness [11, 33, 34], we also included integrins and keratins into the matrisome gene set to provide a broader coverage of the ECM-related genes.

Conversion of Ensembl IDs to gene names was performed either by using the R package biomaRt (version 2.46.2) [35] or the BioMart online tool from Ensembl site (Ensembl release 104) [36].

### Causal analysis

The causal relationship between MCTs and ECMs was evaluated using a hypernetwork modelling approach as previously described [24, 26]. Briefly, hypernetworks represent network structures where edges define a relationship between nodes (e.g., transcripts) and can be shared by many nodes (Figure A1); this is the definition of “higher order interactions” [23]. In this way, hypernetwork structures are used to delineate complex relationships which connect multiple ‘omic elements. We can determine the connectivity of target elements (e.g., differentially expressed genes) within the whole transcriptome by quantifying their number of shared edges (i.e., correlations or gene-gene interactions). This not only provides a summary of higher order interactions from correlation matrices but also implies functional relationships between strongly associated elements [24].

Here, we used hypernetworks to evaluate the correlations between the MCT and ECM (and ECM-related) transcripts within the stromal-epithelial dataset (GSE93326). The differentially expressed MCTs and ECMs were used to refine the selection of target genes for the hypernetworks. To assess the similarity of the differentially expressed MCTs (n=9) or ECM and ECM-related (n=502) transcripts, we first determined the Pearson correlation coefficients as a distance metric between each of these transcripts and the rest of the transcriptome (n=13,815 and n=13,322 in the case of correlations with MCTs and ECMs, respectively). Values were binarized using ±1 standard deviation (sd) from the mean, such that the direction of the correlations (i.e., negative or positive) was ignored. This formed the matrix *M* where,

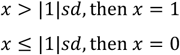

The sd of the r-values were 0.20 and 0.28 for MCT and ECMs, respectively with their distribution profiles shown in Figure A2. The binarized matrix *M* is the incidence matrix of the hypernetwork where target genes are nodes and non-target genes are edges. This matrix was then multiplied by its transpose *M*^*t*^ to generate a new matrix (*M* × *M*^*t*^), which describes the number of shared correlations between any pair of DEGs. This final matrix represents the adjacency matrix of the hypernetwork, which describes the higherorder interactions between the genes, and has been suggested to describe functional relationships [25]. Hierarchical clustering was used to identify the clusters formed within the hypernetwork. The correlations between the highly connected cluster of transcripts from the hypernetwork and the rest of the transcriptome were determined by interrogating the incidence matrix, *M* [26]. This produced a subset of the whole transcriptome which correlated with 90% of the transcripts from the hypernetwork cluster.

The robustness of the hypernetworks was evaluated using a number of approaches. Firstly, we tested if the connectivity of MCT and ECM genes was greater than expected by random chance. This was achieved by iterating one thousand hypernetworks using 7 and 376 randomly selected genes for MCTs and ECMs, respectively, based on the findings of the hypernetwork analysis (Table 2). The mean and sd of the number of ECM and MCT genes that were found to correlate with these random genes were calculated. Secondly, we refined a correlation matrix of both MCTs and ECMs by silencing the indirect relationships between genes, and thus retaining only the direct links [37]. To demonstrate the presence of direct links between the MCTs and ECMs, we first correlated differentially expressed MCT and ECM genes (9 and 502, respectively), together. Indirect relationships in this correlation matrix were silenced using a modified version of equation 5 of Barzel and Barbasi (2013) [37] which incorporated the Moore-Penrose approximation of the inverse of the correlation matrix. Consequently, the matrix *S* represents the directness of the relationship between node (gene) pairs.

All analyses were performed in R (version 4.0.3). The resultant silenced matrix was visualised in Cytoscape (version 3.9.0), imported using the aMatReader app (version 1.2.0). The interaction directness was ranked in Cytoscape to identify the strongest interactions (both positive and negative) between connected nodes. The significance of these interactions was calculated as a z-score (Relationship directness score - mean(S))/sd(S)) and the associated p-values calculated from the cumulative distribution function of the normal distribution.

### Functional annotation

Functional annotation of the genes which showed correlation with the clustered MCT and ECM genes in the hypernetwork analysis was conducted using the online tool DAVID [38, 39].

### Investigation of the correlation between MCT-ECM expression in stroma and epithelium with age at PDAC diagnosis

To investigate how the levels of differentially expressed MCT, ECM and ECM-related genes in stromal and epithelial tissue samples associate with age at diagnosis of PDAC, we performed Pearson correlation analysis using the rcorr function of the R package Hmisc (version 4.5-0; Hmisc: Harrell Miscellaneous (uib.no)). Log-normalised TMM data for 187 samples from the GSE93326 dataset with available age information were used in the correlations. Both the correlation coefficients and the p-values were reported.

### Assessment of MCT, ECM, and ECM-related gene expression in short- and long-term survivors

We evaluated the levels of MCT, ECM, and ECM-related genes identified as differentially expressed in stromal-epithelial dataset in short- and long-term survivors from the GSE79668 dataset (14 and 13 subjects, respectively). Violin plots were used to inspect the log-normalised expression levels of the top four MCT genes which showed differential expression (p<0.05) in stromal-epithelial samples. ECM and ECM-related genes which were significantly different (FDR<0.05) in both stromal-epithelial and STS-LTS datasets were examined with a heatmap using the modified heatmap.2 function as described previously.

## 3. Results

### 3.1. Large inter-study differences in gene expression levels exist in PDAC

In this study, we aimed to establish the link between MCTs and ECMs using publicly available transcriptomic data. Based on the criteria outlined (RNA-seq, primary PDAC tissue biopsies, homo sapiens), four RNA-seq datasets were included in the initial analysis as summarised in Supplementary Table S1 [30, 40-42]. In addition to the studies reporting RNA-seq data from primary PDAC bulk tissues, a study with GEO accession number GSE93326 with data from PDAC stroma and epithelium was selected to help delineate the role of MCTs in desmoplasia. One study was excluded from the analysis due to RNA-seq data being generated from cell lines that were isolated from primary tumours (GSE63124).

The PCA revealed large variation in transcriptomic data from tumour tissues between different datasets whereas minimal differences were observed between tumour and non-tumour tissues (Figure A3). Although the differences between tumour tissues and studies were not clear in the PCA plots when all samples were included in the cluster analysis (Figure A3A-B), this difference was highlighted when tumour stromal and epithelial data were excluded, and the GSE79668 dataset was compared with the remaining datasets (Figure A3C-D). As a result, we deemed the collated RNA-seq datasets on tumour and non-tumour tissue biopsies inappropriate for the purpose of conducting differential gene expression analysis and focused on the analysis of epithelial and stromal data from the GSE93326 dataset.

### 3.2. MCT, ECM, and ECM-related genes are differentially expressed in PDAC stroma and epithelium

The PCA of the GSE93326 dataset (including data from 66 paired epithelium and stroma and additional 57 stroma) showed two distinct clusters of epithelial and stromal samples as similar to previously observed by Maurer *et al*. (2019) for the 60 paired samples [30]. We further investigated whether the separation of epithelial and stromal data was influenced by cancer severity by clustering the samples based on the established PDAC co-variates (which were made available in the GEO database) [43-45]: overall tumour stage, grade, and metastasis to nearby lymph nodes. The distinct separation of stroma and epithelium was not explained by any of the individual factors (i.e., tumour stage) as their occurrence was homogenous among the samples (Figure A4).

Next, we conducted differential gene expression analysis to determine the MCT and ECM genes associated with epithelium and stroma. The analysis showed 13,824 features in 189 samples (123 stroma and 66 epithelium) following filtering of low expression genes. Of these features, 8,944 DEGs were identified between PDAC stroma and epithelium samples (FDR<0.05), including 4,481 upregulated and 4,463 downregulated genes in the stromal compartment (Figure A5, Supplementary Table S2). The genes found to be upregulated in both compartments showed agreement with those found by Maurer *et al*. (2019) [30]. The transcriptional profiles for PDAC stroma were well distinguished from the epithelial counterparts, consistent with the trend observed in PCA (Figure A6).

Nine MCT genes were identified as differentially expressed in PDAC stroma (Table 1, FDR<0.05). Among the MCT genes, *SLC16A10*/MCT10 was the most significant DEG showing nearly 3.5-fold higher expression in stroma relative to epithelium (Figure 1). In addition to the protein coding *SLC16A* genes, we identified a long non-coding RNA (lncRNA) *SLC16A1-AS1* to be differentially expressed and highly upregulated (logFC of 2.25) in stroma samples. Two other *SLC16A* genes that showed higher expression in PDAC stroma were *SLC16A2*/MCT8 and *SLC16A6*/MCT7 with the remaining MCT genes having higher expression levels in epithelium (Figure 1).

**Table 1.**
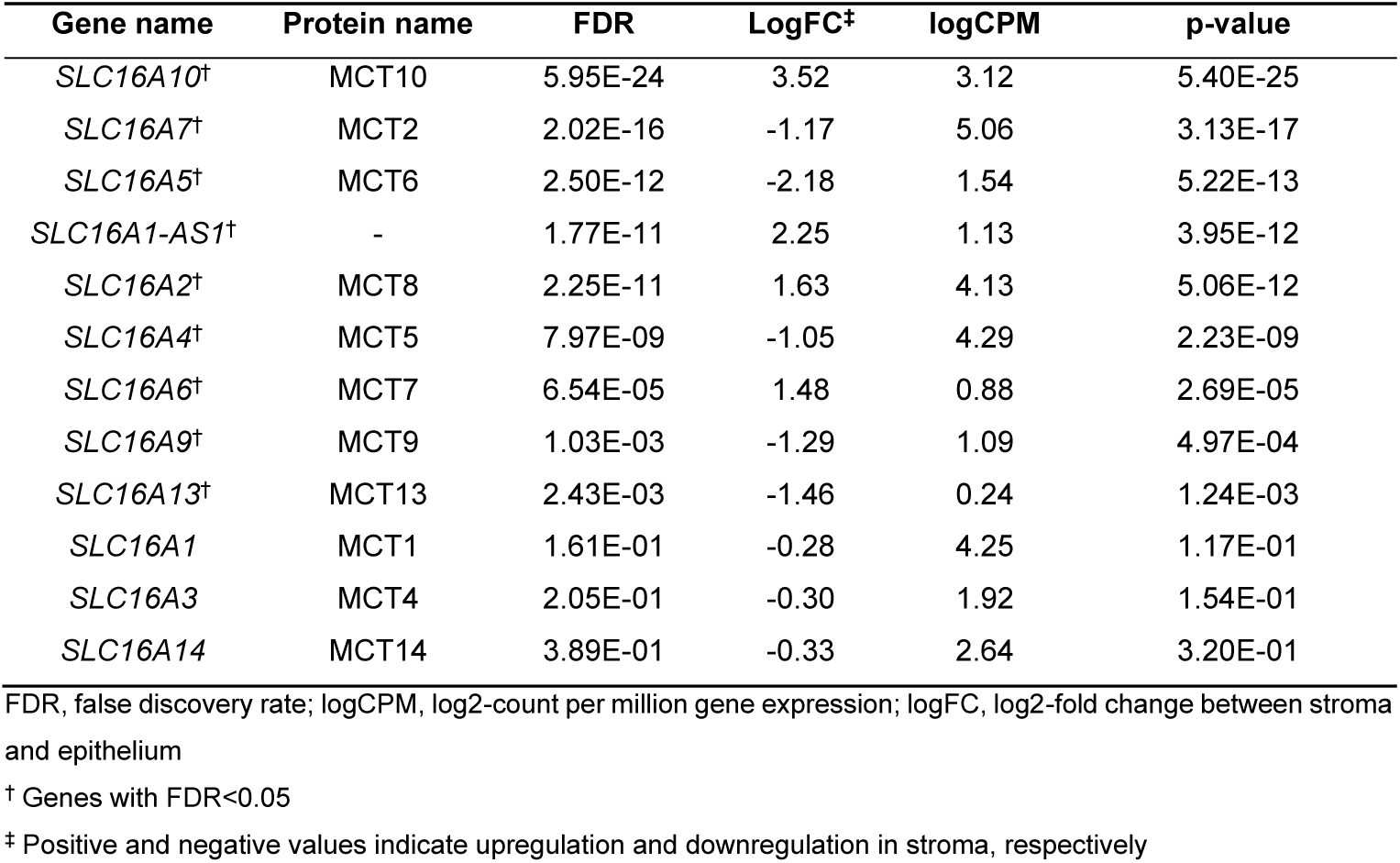
Expression of MCT genes in PDAC stroma (n=123) relative to epithelium (n=66). In total, 12 MCTs were identified, of which, 9 showed significant differential expression between stroma and epithelium with the remaining MCTs having similar expression in these tissues. In addition to the protein coding MCT genes, an lncRNA *SLC16A1-AS1* was also found to be downregulated in tumour epithelium.

**Figure 1.**
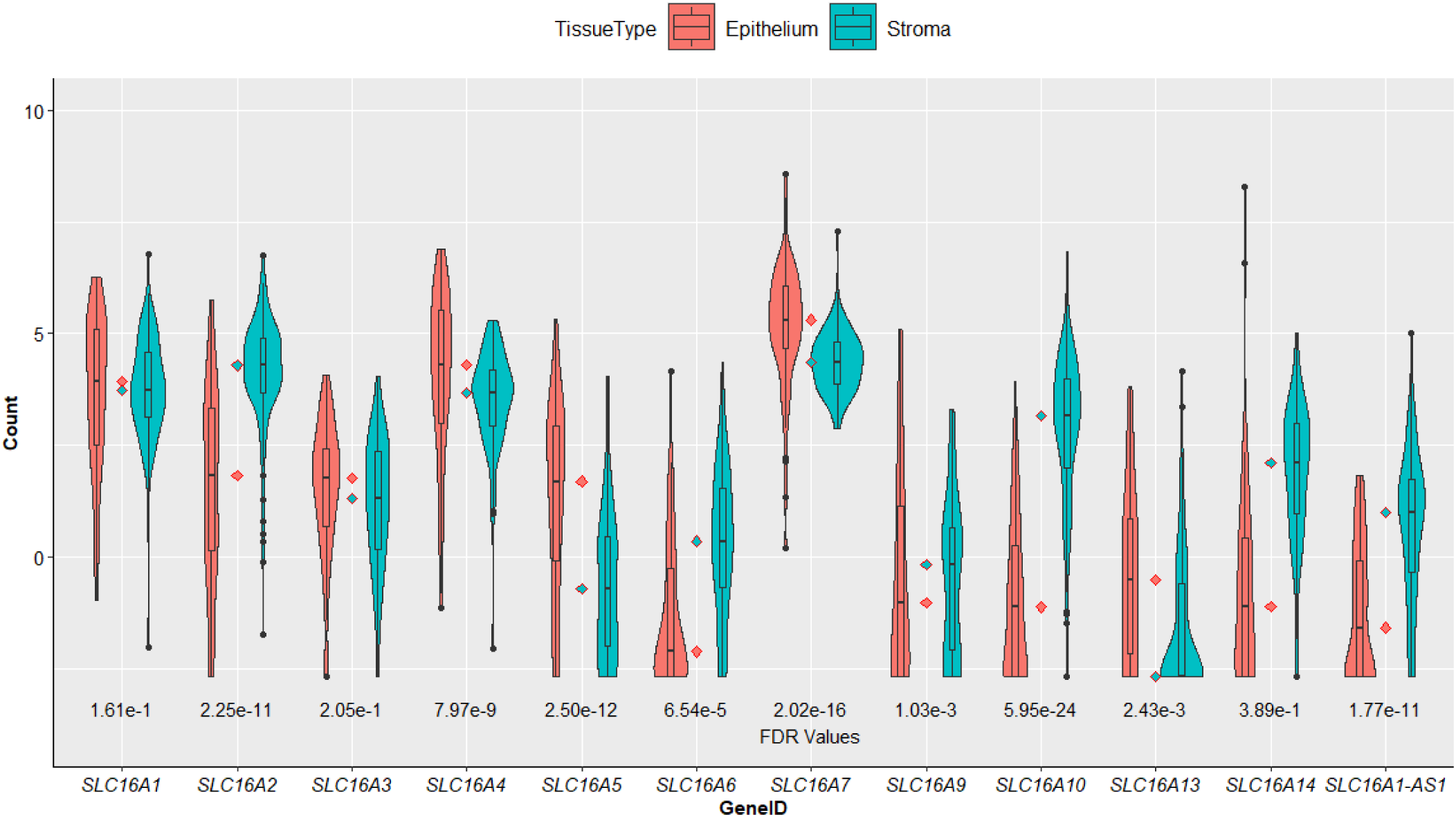
Violin plots of log-normalised expression values of MCT genes in PDAC stroma (n=123) and epithelium (n=66). FDR indicates false discovery rate of 5%. Four MCT genes showed significant upregulation in stroma (*SLC16A2/6/10* and *SLC16A1-*AS1) with 5 MCTs being significantly upregulated in epithelium (*SLC4/5/7/9/13*) and 3 showing no difference in expression between the two tissues. *SLC16A4* and *SLC16A10* showed the largest upregulation in epithelium and stroma, respectively.

To identify differentially expressed ECM and ECM-related genes, we used the NABA_MATRISOME gene set from MSigDB. Using this gene set as a reference and by including integrins and keratins, we identified a total of 502 genes (Supplementary Table S3) including several collagens, laminins, and ECM remodelling enzymes such as lysyl oxidases (*LOX*s) and matrix metalloproteinases (*MMP*s).

### 3.3. There is a causal relationship between MCT and ECM gene expression

Once the differentially expressed MCT, ECM and ECM-related genes in stroma and epithelium samples were identified, a hypernetwork analysis was conducted to assess the relationship between the MCT (n=9) and ECM (n=502) transcripts by revealing the presence of multiple correlations between the groups of genes.

The hypernetwork analysis identified two clusters of MCT transcripts. The first cluster included seven transcripts (*SLC16A1-AS1, SLC16A6, SLC16A5, SLC16A10, SLC16A2, SLC16A13*, and *SLC16A7*) which shared >1500 correlations with the rest of the transcriptome, suggesting similarity and strong functional associations between these set of MCT mRNAs (Figure 2). The second cluster had two MCT transcripts including *SLC16A4* and *SLC16A9* that share <1000 correlations with the rest of the transcriptome, therefore having weaker functional associations with one another and the remaining MCTs.

**Figure 2.**
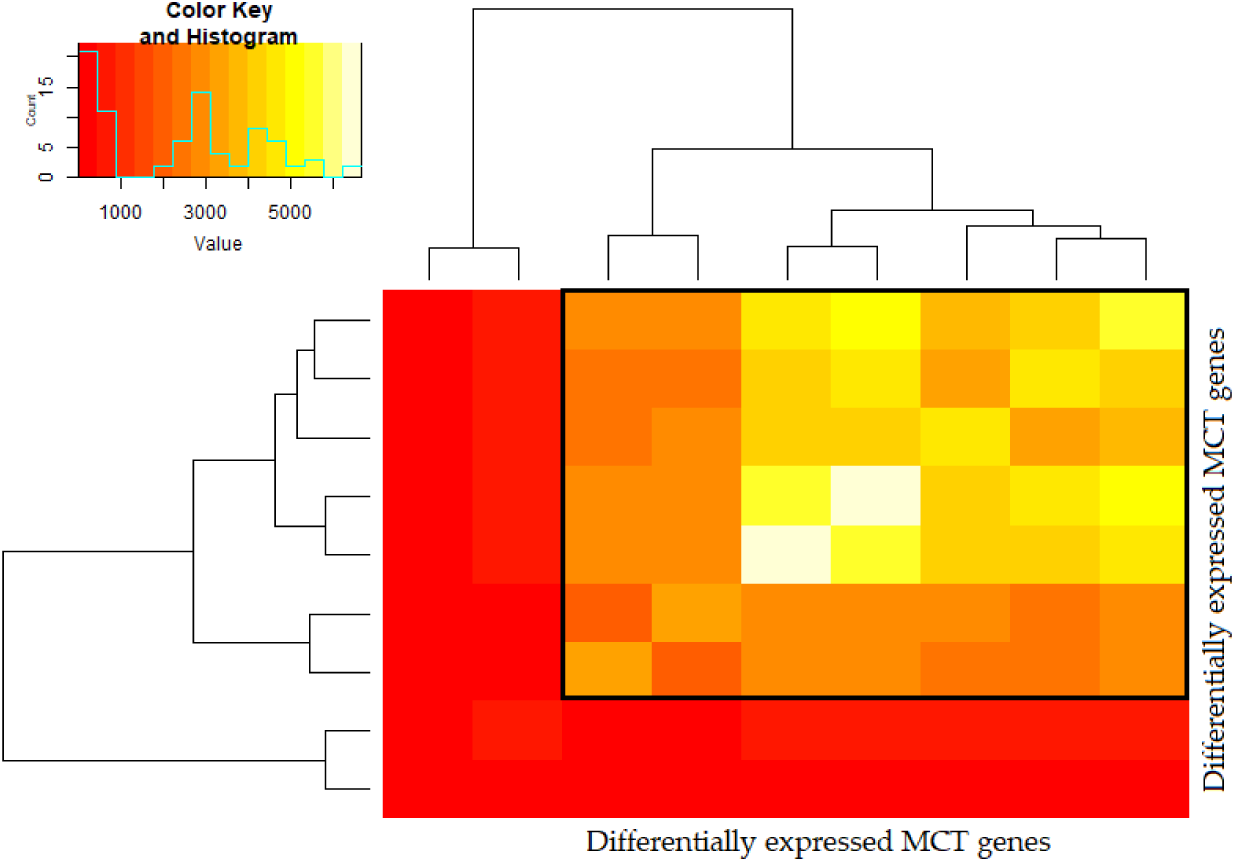
Hypernetwork heatmap of MCT transcripts (n=9) in PDAC stroma and epithelium. Colour intensity represents number of binary relationships with the rest of the transcriptome (n=13,815 total transcripts) shared between a pair of MCT transcripts. Two clusters were identified with cluster 1 having seven MCTs including *SLC16A1-AS1, SLC16A6, SLC16A5, SLC16A10, SLC16A2, SLC16A13*, and *SLC16A7* (highlighted in black box) and cluster 2 having two MCTs including *SLC16A9* and *SLC16A4*.

For the hypernetwork drawn for ECM and ECM-related transcripts, again two clusters were identified including a cluster of 376 transcripts sharing approximately 1000-5000 correlations, and a second cluster of 126 transcripts sharing relatively fewer correlations with the rest of the transcriptome (Figure A7).

Once we identified the functionally associated MCT and ECM transcripts that showed high connectivity with the rest of the transcriptome, we extracted the genes out from the hypernetwork incidence matrix which showed strong correlation with each set of transcripts. We found 1,714 and 2,790 transcripts which correlated with 90% of the 7 MCT and 376 ECM transcripts, respectively. Among the transcripts associated with MCTs, 255 of them were ECM-related including several collagens (*COL1A1/2, COL3A1, COL4A1, COL5A1*), laminins (*LAMA2/3/4, LAMB2, LAMC1/2*), fibronectin 1 (*FN1*) and the ECM remodelling enzymes lysyl oxidase (*LOX*), lysyl oxidase like 1/2/3 (*LOXL1/2/3*), and matrix metalloproteinases (*MMP2/7/11/16/19*) (Table 2, Supplementary Table S4). Likewise, when the hypernetwork analysis was run for ECMs, 4 MCTs were found to associate with them (Table 2). Of the 4 MCTs that associated with ECM and their related transcripts, *SLC16A2*/MCT8, *SLC16A10*/MCT10, and *SLC16A1-AS1* were differentially expressed in stromal-epithelial samples. Likewise, nearly all ECM and ECM-related genes that correlated with MCTs showed differential expression (n=254, Supplementary Table S5). By examining the link between MCTs and ECMs from both directions (i.e., from MCT to ECM transcripts and vice versa) and establishing the bidirectionality of this relationship, we provide evidence for a causal link existing between these sets of genes.

**Table 2.**
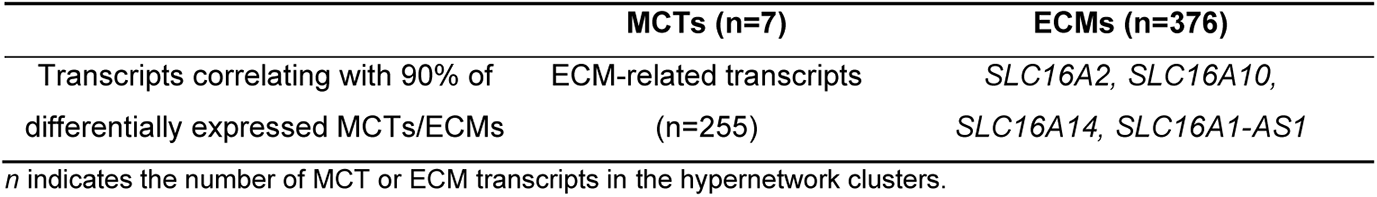
The number of MCT and ECM-related transcripts that showed association with ECM and MCT transcripts in hypernetwork analysis, respectively.

|  | MCTs (n=7) | ECMs (n=376) |
| --- | --- | --- |
| Transcripts correlating with 90% of differentially expressed MCTs/ECMs | ECM-related transcripts (n=255) | *SLC16A2*, *SLC16A10*,<br>*SLC16A14*, *SLC16A1-AS1* |

*n* indicates the number of MCT or ECM transcripts in the hypernetwork clusters.

To confirm that the number of observed interactions between MCTs and ECMs was higher than expected by chance, hypernetworks were iterated using random genes. In this way, the number of MCTs or ECMs associated with a random selection of genes was calculated. Association was defined as previously, where genes must be correlated with 90% of the target genes. When sampling 376 random transcripts representative of the ECMs (Table 2), we found on average 2.45 ± 1.58 MCTs which associated with 90% of the random transcripts after 1000 iterations. When sampling seven random genes, representative of the MCTs, we could not achieve our target of 1000 iterations as we consistently found no genes to be correlated with 90% of our 7 random genes (Table 3). This demonstrated that MCTs were more closely associated with ECMs than the random genes to such an extent that unrelated genes could not replicate our methodology. With this approach the bidirectional causal link between MCTs and ECMs was validated and demonstrated as independent of random chance, however we also used an alternative approach to verify our results.

**Table 3.** Assessment of the robustness of the causal relationship between MCT and ECMs by random sampling of genes 1000 times. The table shows number of MCT and ECM-related transcripts that showed correlation with randomly sampled transcripts given the maximum number of iterations achieved.

|  | Random transcripts<br>representative of MCTs<br>(n=7) | Random transcripts<br>representative of ECMs<br>(n=376) |
| --- | --- | --- |
| Maximum number of iterations achieved | 23 | 1000 |
| Transcripts correlating with 90% of the<br>random transcripts (mean ± SD) | n/a | 2.45 ± 1.58 |
| n/a, not available |  |  |

Analysis of the direct paths in a correlation matrix containing MCT and ECM genes resulted in a silenced network *S* consisting of 511 nodes (genes) and 130,305 edges (gene-gene interactions). Having ranked directness between node pairs, we found that some interactions (both positive and negative) between ECMs and MCTs were among the strongest interactions in the network. Included in this selection were *SLC16A2*/MCT8, *SLC16A1*-AS1, and most commonly, *SLC16A10*/MCT10 which were all among the most highly connected MCT genes in our hypernetwork (Table 4). This approach provides further validation, independent of the hypernetworks, for a causal link between MCTs and ECMs.

**Table 4.**
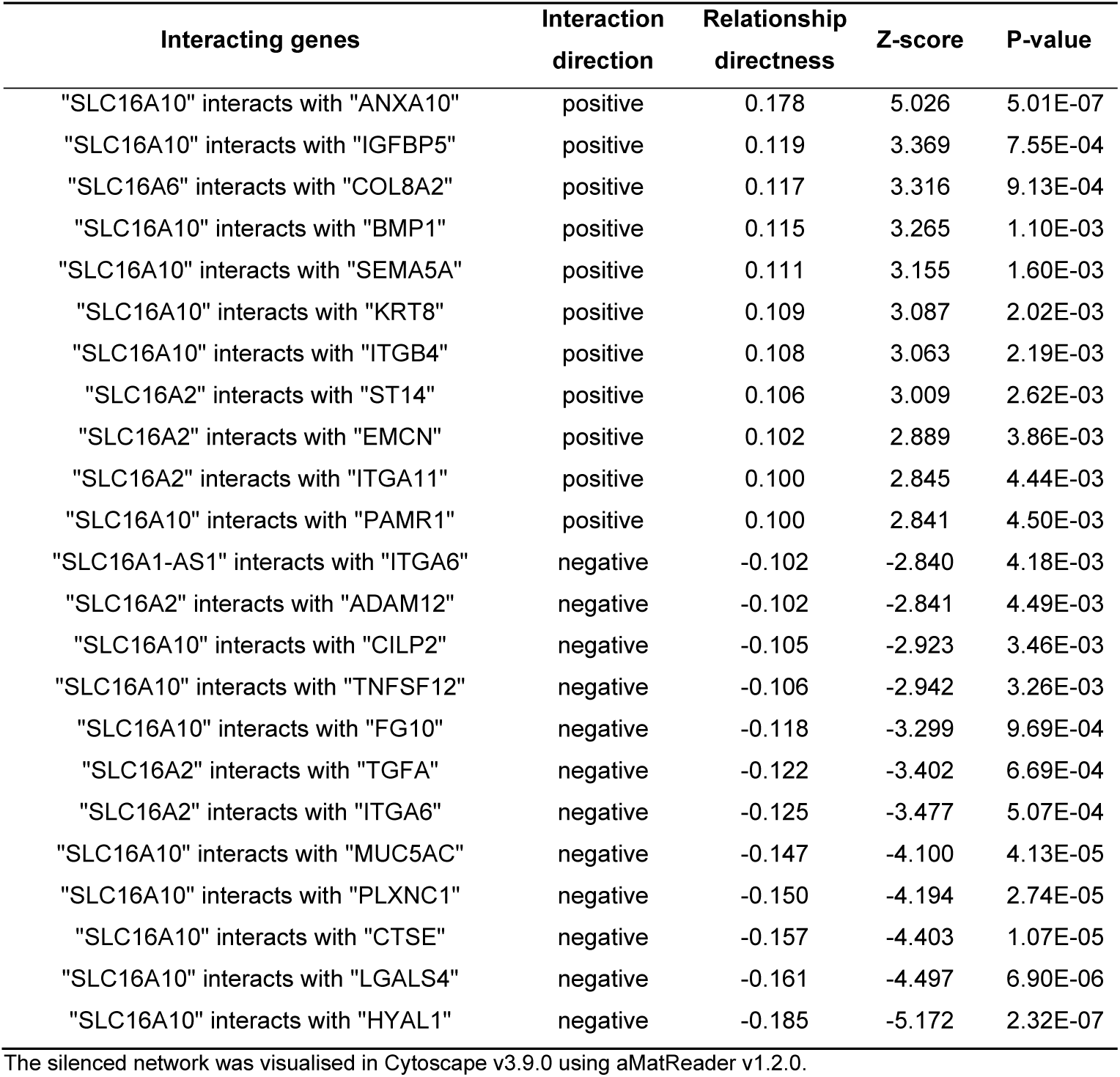
The strongest 23 interactions found between MCT and ECM genes within the silenced network. Values from the matrix *S* describing the directness of the relationship between node pairs are presented alongside a z-score describing the position of this value in the whole network and the associated p-value of this position. A directness value of 0.1 was used as it is approximately a z-score of 3 (97.5% in a two-tailed test). All interactions presented below are statistically significant.

### 3.4. Lactate and thyroid hormone transporters correlate with ECMs involved in cancer associated signalling pathways

Functional annotation of MCT (n=7) and ECM (n=254, differentially expressed and correlated with MCTs) transcripts using DAVID revealed the presence of ECM components involved in ECM-receptor interaction, protein digestion and absorption, phosphoinositide-3-kinase–protein kinase B (PI3K-Akt) signalling pathway, mitogen-activated protein kinase signalling pathway, and pathways in cancer (Supplementary Table S6). Extracellular matrix organisation and disassembly, cell adhesion, collagen catabolic process, cell adhesion by integrin, and plasma membrane lactate transport were found as biological processes associated with ECMs and MCTs, respectively. In addition to lactate transport as a biological process for most MCTs, thyroid hormone transport was highlighted for *SLC16A2*/MCT8 and *SLC16A10*/MCT10.

### 3.5. MCT and ECM mRNA levels are not associated with age at PDAC diagnosis

We investigated if the levels of MCT, ECM, and ECM-related mRNAs in stromal and epithelial samples (n=187) associated with age at PDAC diagnosis by a Pearson correlation analysis. The age at diagnosis ranged between 38–89 with mean and median of 68.3 and 69, respectively.

We found that there was no strong correlation between the levels of any of these mRNA transcripts with age at diagnosis (Supplementary Table S7 and Table S8).

### 3.6. SLC16A3/MCT4 and several ECM components are significantly upregulated in STS subjects

When we examined the expression of the top four MCT genes including *SLC16A5*/MCT6, *SLC16A7*/MCT2, *SLC16A10*/MCT10, and *SLC16A1-AS1* in stromal-epithelial samples in short-and longterm survivors of PDAC (14 and 13 subjects, respectively) from the GSE79668 dataset, we found that none of these MCT genes showed differential expression in these subjects (FDR>0.05). The only MCT gene that was significantly upregulated in STS subjects was *SLC16A3*/MCT4 (FDR=0.02, logFC=1.55). However, the unadjusted p-values of these MCTs indicated significant upregulation of *SLC16A7* (logFC=1.3) and downregulation of *SLC16A1-AS1* (logFC=-1.0) in STS subjects (p-values of 0.003 and 0.006, respectively) (Figure 3).

**Figure 3.**
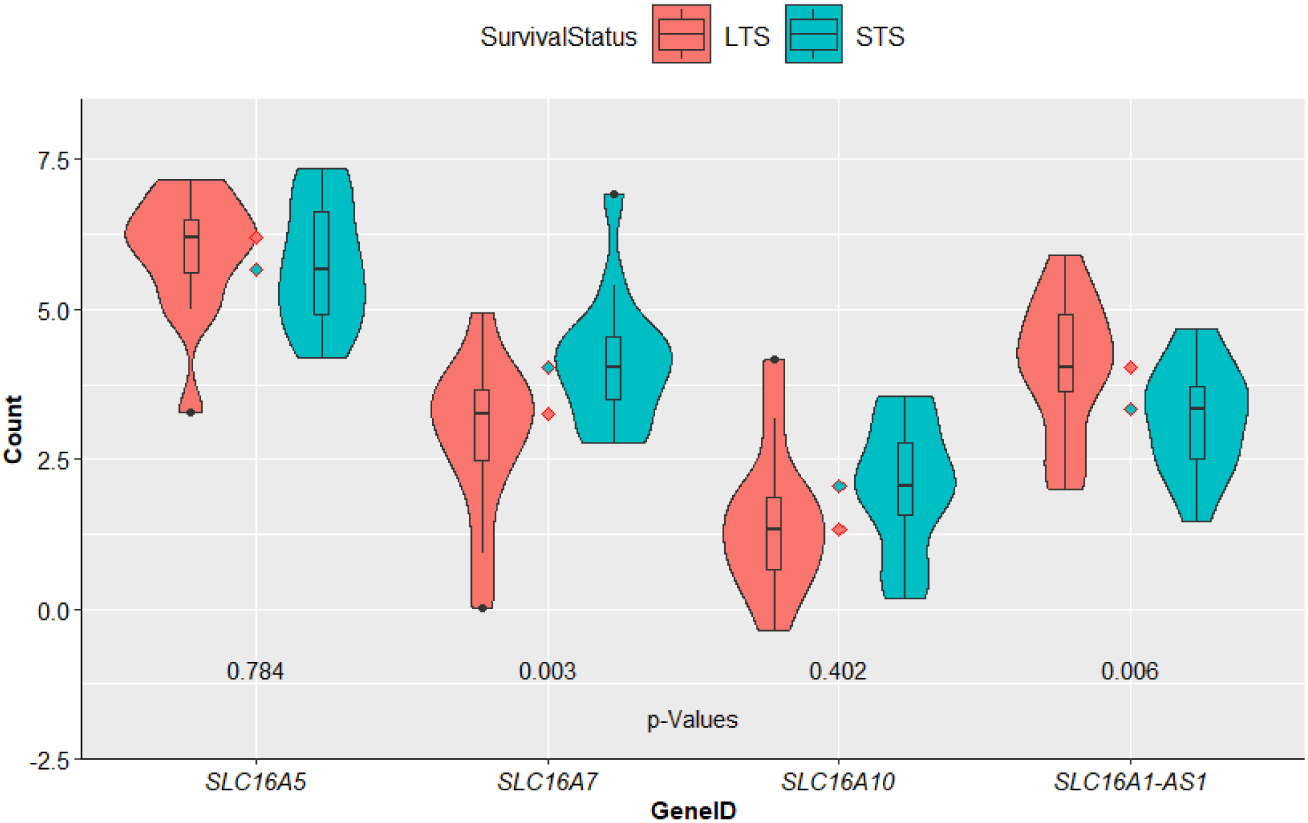
Log-normalised levels of MCT transcripts *SLC16A5*/MCT6, *SLC16A7*/MCT2, *SLC16A10*/MCT10, and *SLC16A1-AS1* in short- and long-term survivors (STS and LTS, respectively) from the GSE79668 dataset. These four genes are the top 4 differentially expressed MCTs in PDAC stroma-epithelium (FDR<0.05) from the GSE93326 dataset, hence the rationale for selecting them for investigation in the GSE79668 dataset. The number of samples from STS and LTS subjects are 14 and 13, respectively. Indicated p-values are not FDR-adjusted. Of the four MCTs investigated, *SLC16A7* and *SLC16A1-AS1* were significantly upregulated in STS and LTS subjects, respectively with other two MCTs showing similar expression in both survival groups (p<0.05).

Among the ECM and ECM-related genes that were differentially expressed in stromal-epithelial samples (n=255), only 15 of them were significantly expressed with majority being upregulated in the STS subjects (Figure 4, FDR<0.05) including *CTSH, S100A2, SERPINE1, WNT7B, GPC3* and a number integrins (*ITGA3* and *ITGA6*).

**Figure 4.**
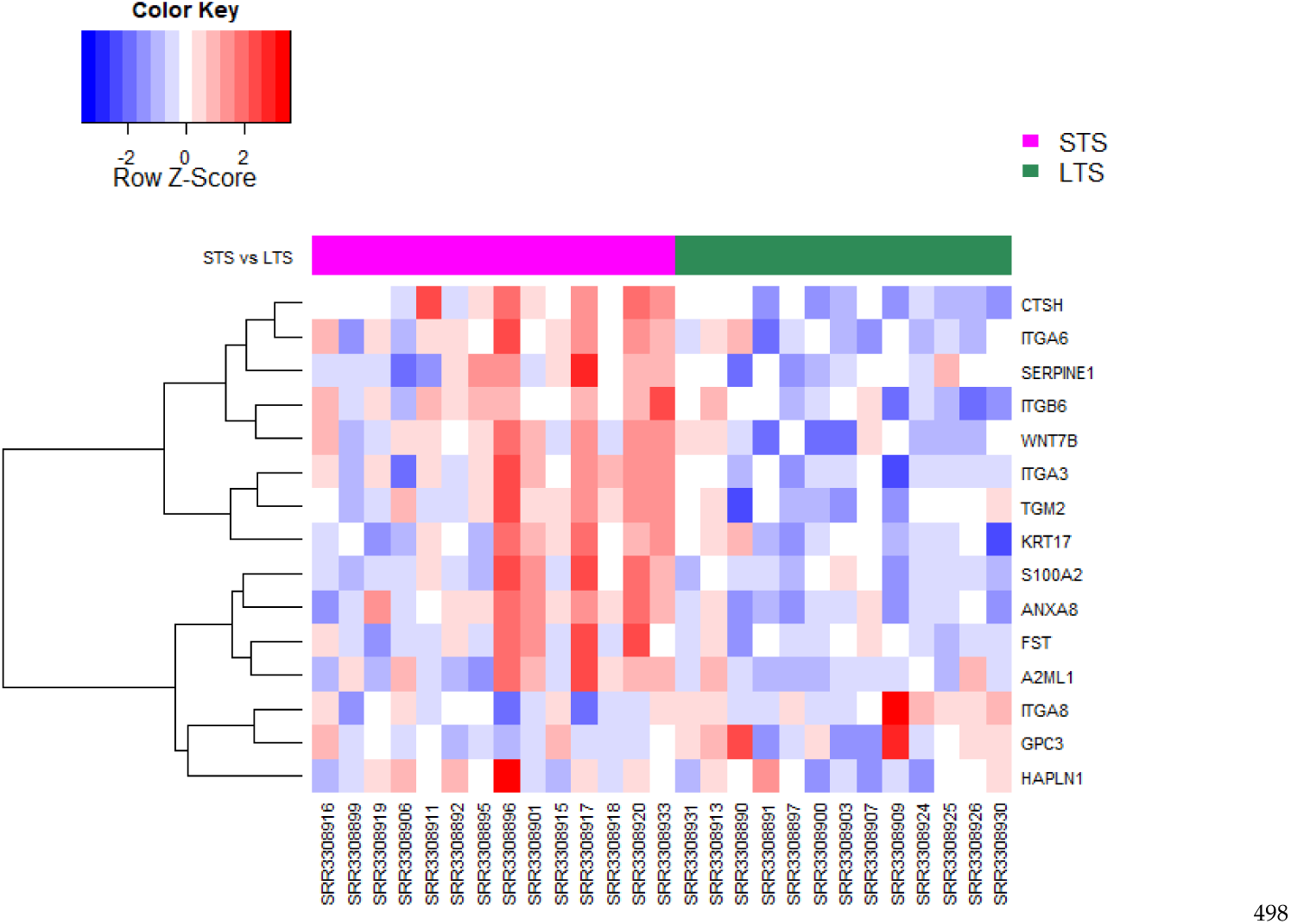
Heatmap and dendrogram showing the normalised expression levels of ECM-related genes (n=15) which were differentially expressed in PDAC stroma-epithelium as well as in short-and long-term PDAC survivors (STS and LTS, respectively) from the dataset GSE79668 (FDR<0.05). The number of samples from STS and LTS subjects are 14 and 13, respectively. Red and blue colours indicate increased and decreased expression, respectively.

## 4. Discussion

PDAC is a devastating cancer with high morbidity and low survival rates worldwide with no effective medical treatments available to improve patient prognosis. The drawbacks of existing treatment strategies for PDAC and lack of improvement in patient outcomes necessitate the need for a better understanding of tumour microenvironment and the mechanisms contributing to disease progression [5, 7, 46]. In this study, we hypothesized that MCTs contribute to PDAC desmoplasia by associating with ECM components. We sought to explore the presence of a causal relationship between MCTs and ECMs to suggest that the increased expression of ECM components may be linked with increased MCT expression.

Our first aim was to understand the expression levels of MCT mRNAs in PDAC biopsies relative to nontumour tissues as such a comparison has not been widely explored using RNA-seq data. The unsupervised and hierarchical clustering analysis of the collated data from GEO showed no separation of the tumour and non-tumour samples indicating the similarity of these tissue types. In addition, the hierarchical clustering of tumour and non-tumour samples indicated large inter-study variation in gene expression. The large inter-tumoral variation may perhaps be explained by the highly heterogenic nature of PDAC disease and consequently the biopsy samples which was also highlighted in previous studies [4, 30]. In addition to biological variation, technical factors such as procedures used in tissue acquisition, the sequencing platform, and library preparation method may also contribute to such heterogeneity. Even with the use of same sample acquisition technique, differences in other mentioned factors may still lead to variation in the data. For example, close examination of the tumour cluster in Figure A3A-B shows some separation of the tumour samples from the GSE93326 and GSE79668 datasets although both studies employed macrodissection of the bulk tissue following surgical resection [30, 40]. It was also interesting to see the lack of separation between the tumour and non-tumour samples which may be influenced by the purity of bulk tissue during sample collection. Another factor which may have played a role could be the lack of properly paired samples as we only had a single dataset which included paired tumour and non-tumour tissues with limited sample size with remaining datasets not including non-tumour tissues. As a result of the large patient variability and the limited size of non-tumour samples, we concluded that conducting differential gene expression analysis on tumour and non-tumour samples using the combined dataset would be inappropriate.

We subsequently focused on the stromal and epithelial dataset. This is because MCTs are reported in both stroma and epithelium and in PDAC, the main ECM production happens in stromal cells. Hence it is important to understand and correlate expression of MCTs and ECMs in different compartments. Our analysis revealed nine differentially expressed MCT genes in both stromal and epithelial samples including the upregulated thyroid hormone (TH) transporters MCT8 (*SLC16A12*) and MCT10 (*SLC16A10*), monocarboxylate transporter MCT2 (*SLC16A7*), MCT6 (*SLC16A5*) with potential role in glucose and lipid metabolism, and an lncRNA *SLC16A1-AS1* previously identified as a prognostic or diagnostic biomarker in a number of cancers [13, 47-54]. To our knowledge, the upregulation of the TH transporters in the stromal-epithelial samples has not been highlighted previously. TH transporters regulate the availability of THs T_3_ and T_4_ in cells based on their levels in the local tissue. THs themselves are key regulators of energy metabolism, growth, differentiation, and physiological function of tissues [47, 55]. The role of THs in cancer has been pointed in several studies and their association been summarised in a recent review [56]. For example, induction of cell proliferation and metabolism by T_3_ via the activation of TRβ1/Akt pathway was demonstrated in human pancreatic insulinoma cells [57]. In addition, inhibition of TH binding to their cell surface receptor integrin αVβ3 was shown to inhibit tumour cell proliferation and angiogenesis in *in vitro* and *in vivo* xenograft models of pancreatic cancer [58]. A more recent study proposed pharmacologically induced TH inactivation as a strategy to reduce tumour metastatic potential based on their observation that T_3_ promoted epithelial-mesenchymal transition by transcriptional activation of ZEB-1, mesenchymal genes and MMPs and suppression of E-cadherin, thereby influencing tumour progression and metastasis in skin squamous cell carcinoma [59]. Furthermore, a link between THs and ECM organisation has been shown where T_3_ was suggested to induce secretion of growth factors that stimulate cellular proliferation, ECM reorganisation involving fibronectin and laminin, and changes in cell spreading and adhesion [60]. In contrast to the influence of THs on tumour progression discussed above, the synergistic action of T3 in combination with gemcitabine and cisplatin was shown to enhance the cytotoxicity of the chemotherapy agents in *in vitro* models of pancreatic cancer, highlighting the benefit of the combination therapy for the treatment of pancreatic cancer [61]. Despite the presence of associations between thyroid dysfunction including hypothyroidism and increased the risk of pancreatic cancer, the effects of THs in cancer neoplasia is currently conflicting [47, 56, 62]. Therefore, better understanding of the molecular mechanisms involved in contribution of THs to pancreatic cancer is needed. The lncRNA *SLC16A1-AS1* has previously been found to be upregulated in osteosarcoma, glioblastoma and oral squamous cell carcinoma [50, 52, 63], downregulated in non-small cell lung cancer and cervical squamous cell carcinoma [19, 49], and show conflicting expression profile in hepatocellular carcinoma [51, 53, 54]. This is the first study highlighting differential expression of *SLC16A1-AS1* in PDAC showing lower levels of expression in epithelial relative to stromal compartment. With increasing number of reports showing *SLC16A1-AS1* as a potential biomarker in cancer, we propose an investigation to understand how the upregulation of this lncRNA in PDAC associates with tumour development, progression and overall survival.

Our second primary goal was to explore the causal relationship between MCTs and ECMs in PDAC and understand if the expression of the latter is influenced by upregulation of MCTs. We used hypernetworks to investigate the relationships between MCTs and ECMs and tease out the functional connectivity among the individual elements. Our analysis indicated 7 MCTs with functional association in a pairwise manner including the lncRNA *SLC16A1-AS1*, MCT7 (*SLC16A6*), MCT6 (*SLC16A5*), MCT10 (*SLC16A10*), MCT8 (*SLC16A2*), MCT13 (*SLC16A13*), and MCT2 (*SLC16A7*). The extraction of the genes associated with MCTs from the hypernetwork matrix revealed 255 ECMs which showed correlation with 90% of these MCTs including several collagens, laminins, fibronectin 1 and the ECM remodelling enzymes. Similar analysis conducted for ECMs indicated 376 ECM transcripts which were functionally connective and correlated with 4 MCTs including the differentially expressed *SLC16A1-AS1, SLC16A2*/MCT8 and *SLC16A10*/MCT10. Hypernetworks generated for both MCTs and ECMs demonstrate the presence of bidirectional associations and suggest a causal link exists between these two sets of transcripts. To rule out the randomness of this observation, we conducted a robustness testing by firstly generating hypernetworks with random transcripts that were of identical size to the number of MCTs and ECMs which were clustered in our analysis. The difficulty we experienced in executing iterations with random genes in replacement of MCTs showed that relationships between MCTs and ECMs were stronger than those which were frequent between random genes (Table 3). When the analysis was repeated for ECMs, we found 2.45 ± 1.58 MCTs which were associated with ECMs, similar to our primary finding. As a result, we further conducted a direct path analysis which aimed to refine the correlation matrix of MCTs and ECMs and reveal only the direct gene-gene interactions with causal relationship by silencing indirect connections. Correlations in biological networks that retain both direct and indirect links can confound identification of true pairwise interactions [37]. Independent from the hypernetwork modelling approach used, the direct path analysis confirmed the presence of significant MCT-ECM interactions, involving particularly *SLC16A10*/MCT10, which were among the strongest of all interactions observed within the whole silenced network. This analysis therefore further reinforces that the connectivity of MCTs and ECMs is not by random chance and the causal relationship between the MCT and ECM action is robust. The mechanisms of the direct interactions between the MCTs and ECMs identified in this study is currently unknown and requires further elucidation. One potential facilitator of this interaction may be CD147, an MCT chaperone and regulator, which is a key contributor to tumour growth and metastasis by promoting ECM remodelling through the induction of MMPs [64]. The expression of CD147 is influenced by its association with MCTs and the synergistic action of both proteins could enhance metastasis via acidification of the tumour microenvironment and degradation of ECM by MMPs [17].

The tumour biomarkers are undoubtedly valuable in several aspects of patient management including screening, diagnosis, monitoring, or patient stratification for an intended cancer therapy. Given the insidious nature and poor prognosis of PDAC, identification of novel biomarkers for early detection, personalised therapy and post-resection follow-up are urgently needed to improve patient overall survival. As ECM is a driver of tumorigenesis, use of ECM-derived biomarkers can immensely facilitate diagnosis and prognosis of patients with cancer in the clinic [65]. To this end, we evaluated the presence of correlations between the levels of MCT and ECM components in stromal and epithelial samples and age at PDAC diagnosis. Many of the ECM components that showed differential expression and were highlighted in the hypernetwork analysis, including *MMPs, TIMP3, FN1, LAMC1*, and *COL4A1/6A2*, have previously been reported as prognostic or diagnostic markers in different cancers [65]. In our analysis, we found no association between any of the MCTs or ECM components with age at diagnosis. However, scrutinising the expression levels of these ECM components in short-term survivors of PDAC revealed a number of genes which were upregulated in these subjects and have been suggested to have a diagnostic, predictive or prognostic value in different cancers including *S100A2* (pancreatic cancer), cathepsin H/*CTSH* (thyroid carcinoma), *SERPINE1* (gastric cancer, oesophageal cancer), *WNT7B* (breast cancer, colorectal cancer), and *GPC3* (hepatocellular carcinoma) [66-73]. Although the latter analysis using the GSE79668 dataset comes from a rather small dataset and poses a limitation, the findings still highlight some potential biomarkers that could be investigated in PDAC. Finally, of the significant interactions identified in direct path analysis, *SLC16A10*/MCT10 showed the strongest interaction with *ANXA10*, a potential marker associated with the progression of pancreatic precursor lesions to PDAC [74], which opens an avenue for further investigation.

## 5. Conclusions

Our transcriptomic analysis revealed multiple MCTs in PDAC stromal and epithelial compartments including lncRNA *SLC16A1-AS1* which may serve as a novel potential diagnostic or prognostic biomarker. In addition, a subset of the differentially expressed ECM components showed associations with MCTs, such as collagens, laminins, fibronectin 1, and ECM crosslinking and remodelling enzymes, highlighting the role of MCTs in PDAC desmoplasia which should be considered when developing future treatment strategies to improve patient outcomes. Analysis of the higher order interactions through hypernetworks indicates the presence of a causal link between MCTs and ECMs and warrants the need for further studies to elucidate their functional connection.

## Supplementary Materials

Table S1: The summary of the RNA-sequencing studies of pancreatic ductal adenocarcinoma (PDAC) samples collated from GEO, Table S2: List of differentially expressed genes in PDAC stroma relative to epithelium, Table S3: List of differentially expressed ECM and ECM-associated genes in stroma relative to epithelium, Table S4: The identity of the ECM and associated transcripts (n=255) correlating with 90% of the MCTs in the central cluster (n=7), Table S5: The ECM and their related transcripts (n=254) that correlated with MCTs in a bilateral manner, Table S6: Functional annotation of differentially expressed MCT (n=7) and ECM (n=254) genes highlighted by hypernetwork analysis, Table S7: Correlation of MCT transcripts with age (ranging between 36-86) at cancer diagnosis. Pearson correlation analysis for each MCT was conducted for n=187 samples, Table S8: Correlation of ECM and their related transcripts with age (ranging between 36-86) at cancer diagnosis. Pearson correlation analysis was conducted for n=187 samples.

## Author Contributions

Conceptualization, A.L., A.S.; methodology, T.G., A.S.; software, T.G., A.S.; formal analysis, A.U., T.G.; investigation, A.U., T.G.; resources, A.S., A.L.; data curation, A.U.; writing— original draft preparation, A.U.; writing—review and editing, A.U., T.G., A.S., A.L.; visualization, A.U., T.G., A.S., A.L.; supervision, A.S., A.L.; project administration, A.S., A.L.; funding acquisition, A.L.

## Institutional Review Board Statement

Not applicable.

## Data Availability Statement

Publicly available datasets were analyzed in this study. The data can be found in GEO repository: GSE119794, GSE93326, GSE79668, and GSE131050. The source codes are available at GitHub repository: https://github.com/aysheu/pancreatic-cancer-pipeline.

## Appendix A

**Figure A1.**
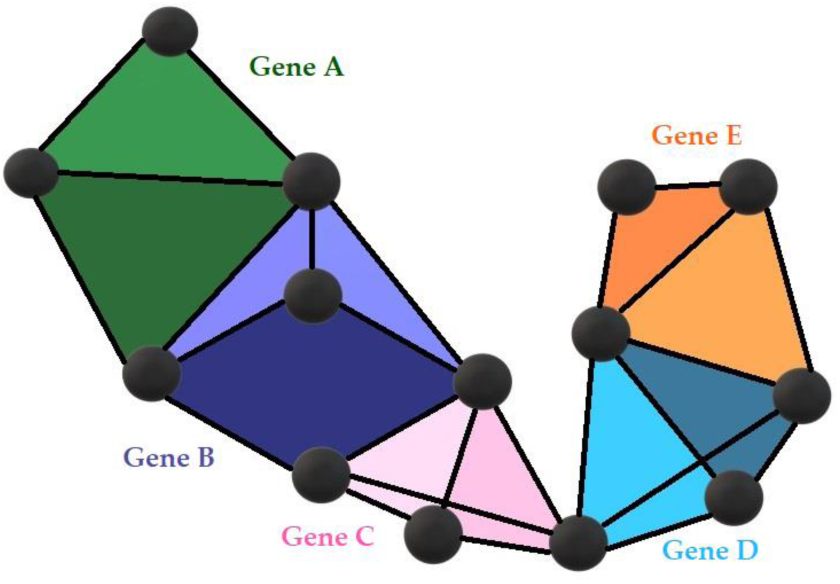
A hypernetwork diagram with three-dimensional structure of genes that correlate with the expression of other genes (black spheres). Shared correlations are shown by the black connecting edges.

**Figure A2.**
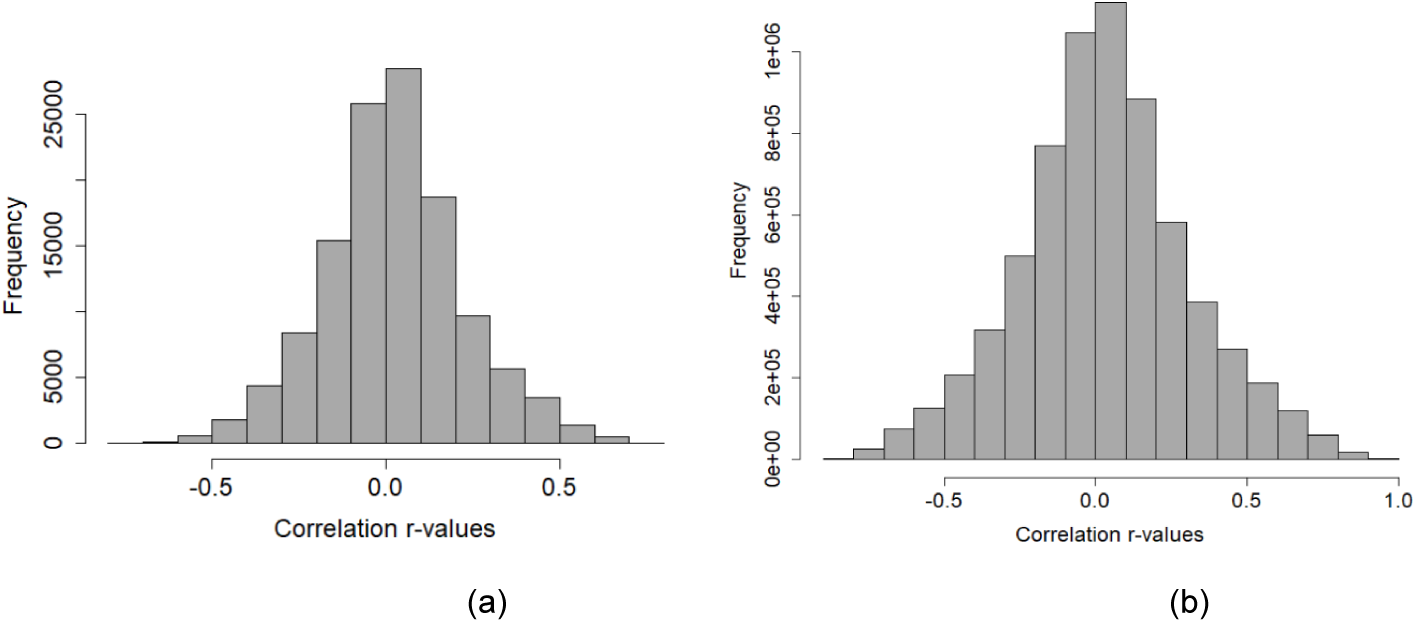
The frequency of the observed coefficient values resulted from the correlation of differentially expressed a) MCTs and b) ECMs with the rest of transcriptome in PDAC stroma.

**Figure A3.**
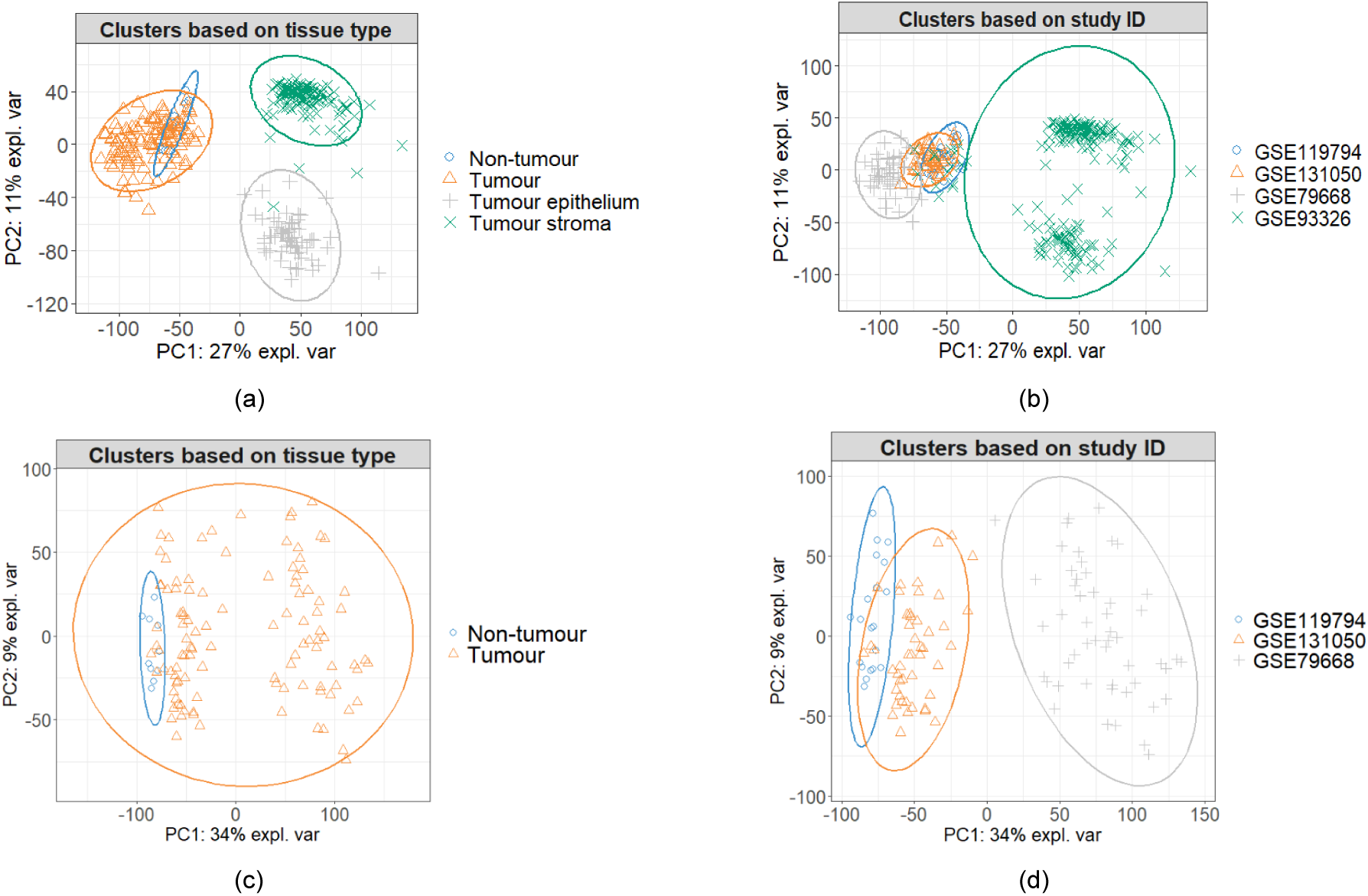
Unsupervised clustering of the collated datasets using Principal Component Analysis. Clustering of a) all datasets based on tissue type (tumour=123, non-tumour=10, tumour epithelium=66, tumour stroma=123), b) all datasets based on GEO accession number, c) all datasets excluding GSE93326 based on tissue type (tumour=108, nontumour=10), and d) all datasets excluding GSE93326 based on the GEO accession number.

**Figure A4.**
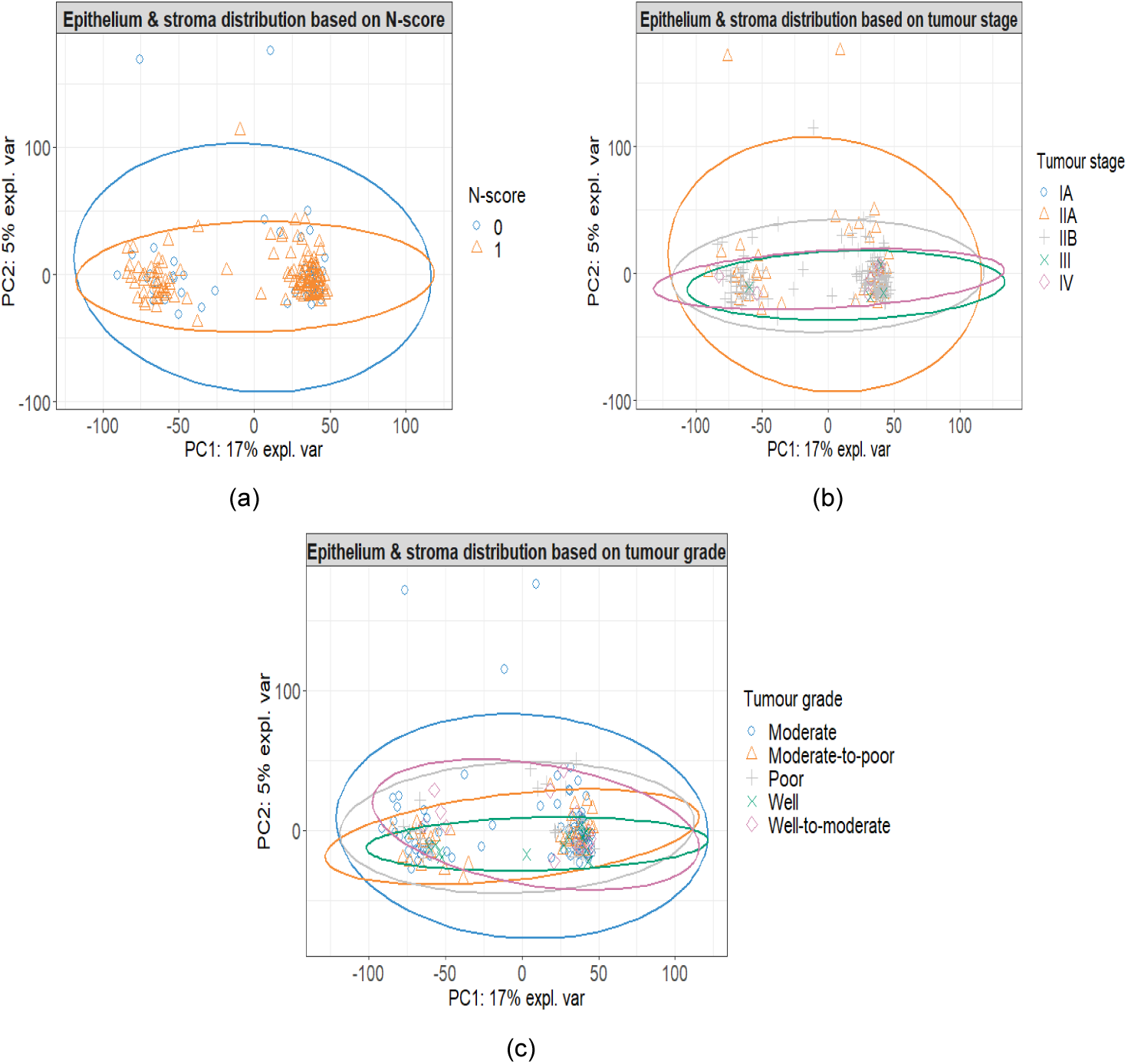
Unsupervised clustering of 187 samples with available metadata (65 epithelium and 122 stroma) from the GSE93326 dataset using Principal Component Analysis. Clustering of samples was performed based on a) tumour N-score (n=177), b) overall tumour stage (n=187), and c) tumour grade (n=187). N-score: 0 and 1 indicate no spread to nearby lymph nodes and spread to no more than three nearby lymph nodes, respectively.

**Figure A5.**
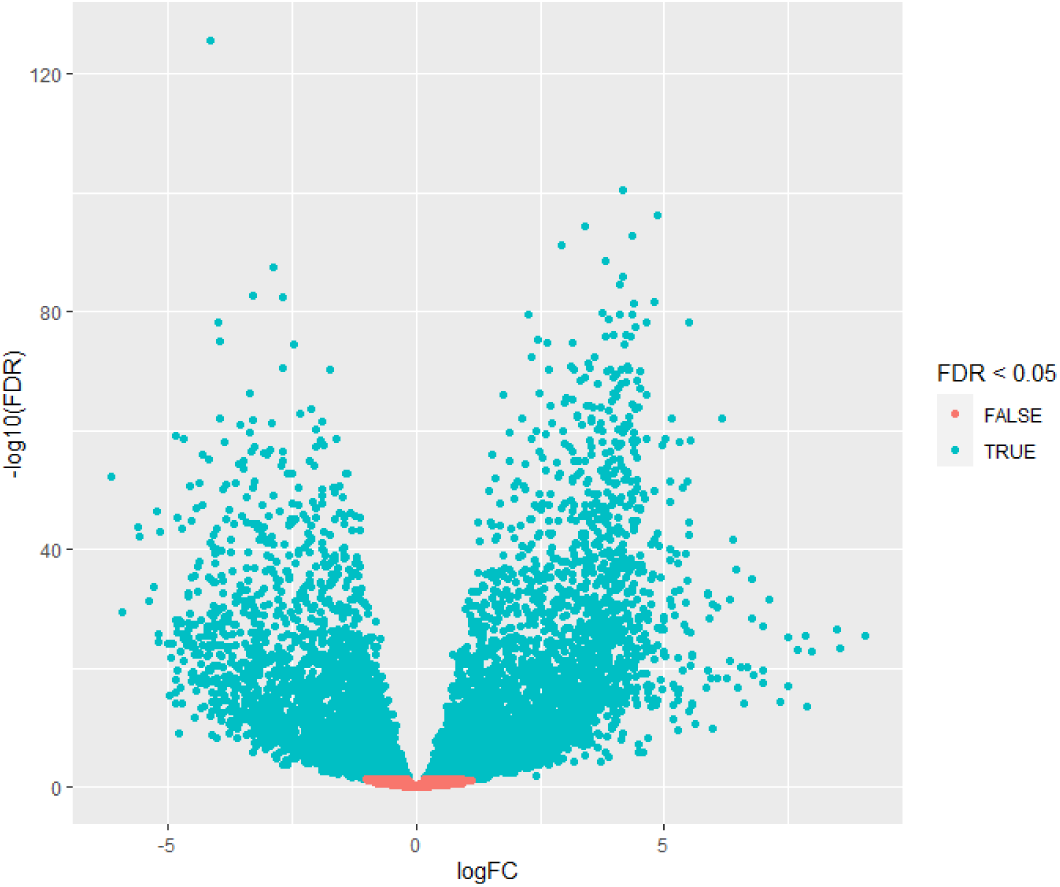
Volcano plot of the genes (n=13,824) after filtered for low expression in PDAC stroma (n=123) and epithelium (n=66) samples. Sixty-six out of 123 stroma samples are matching counterparts of epithelium samples. Cyan and orange dots indicate genes that are differentially expressed (n=8,944) and no change in expression between epithelium and stroma (n=4,880), respectively (FDR<0.05).

**Figure A6.**
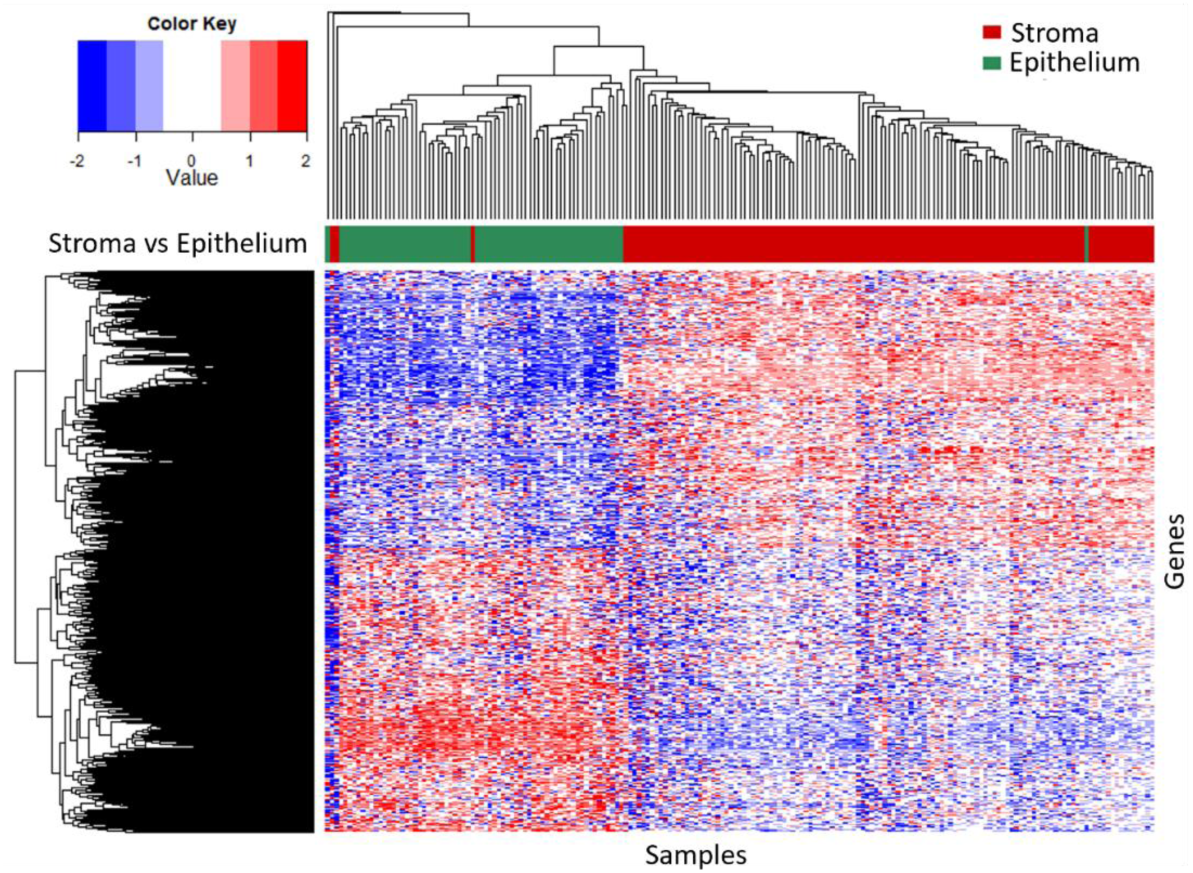
Heatmap and dendrogram showing the normalised and scaled (z-scored) expression levels of genes (n=8,944) differentially expressed in PDAC stroma relative to epithelium (FDR<0.05), across n=189 samples. Red and blue colours indicate increased and decreased expression, respectively.

**Figure A7.**
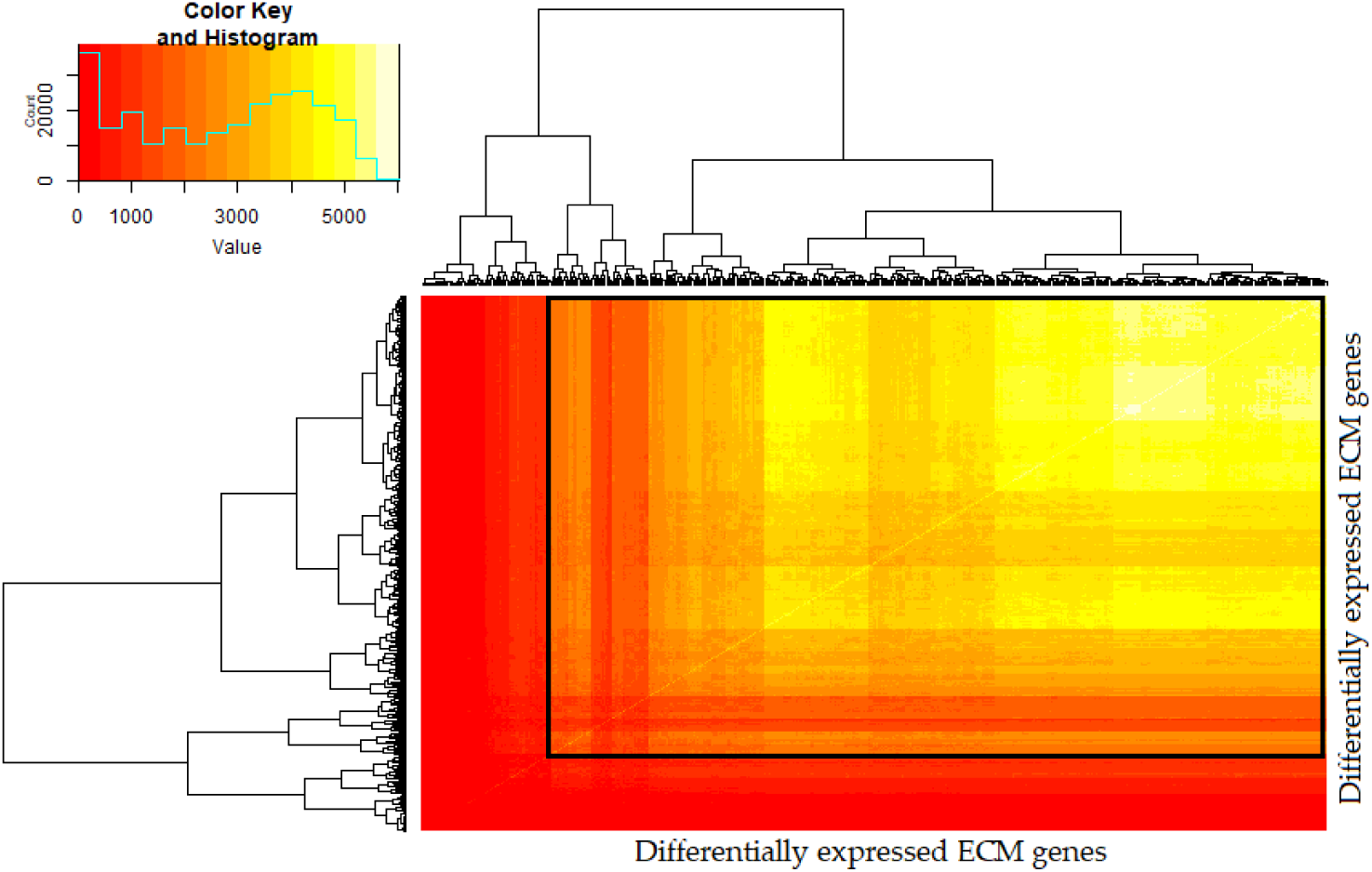
Hypernetwork heatmap of ECMs (n=502 transcripts) in PDAC stroma. Colour intensity represents number of binary relationships shared between a pair of transcripts with the rest of the transcriptome (n=13,322 total transcripts). Two clusters were identified with cluster 1 having 376 ECMs with higher connectivity (highlighted in black box) and cluster 2 having 126 ECMs with relatively fewer correlations with the rest of the transcriptome.

